# Breeding seasonality generates reproductive trade-offs in a long-lived mammal

**DOI:** 10.1101/2021.01.19.427238

**Authors:** Jules Dezeure, Alice Baniel, Alecia J. Carter, Guy Cowlishaw, Bernard Godelle, Elise Huchard

## Abstract

The evolutionary benefits of reproductive seasonality are usually measured by a single fitness component, namely offspring survival to nutritional independence (Bronson, 2009). Yet different fitness components may be maximised by dissimilar birth timings. This may generate fitness trade-offs that could be critical to understanding variation in reproductive timing across individuals, populations and species. Here, we use long-term demographic and behavioural data from wild chacma baboons (*Papio ursinus*) living in a seasonal environment to test the adaptive significance of seasonal variation in birth frequencies. Like humans, baboons are eclectic omnivores, give birth every 1-3 years to a single offspring that develops slowly, and typically breed year-round. We identify two distinct optimal birth timings in the annual cycle, located 4-months apart, which maximize offspring survival or minimize maternal interbirth intervals (IBIs), by respectively matching the annual food peak with late or early weaning. Observed births are the most frequent between these optima, supporting an adaptive trade-off between current and future reproduction. Furthermore, infants born closer to the optimal timing favouring maternal IBIs (instead of offspring survival) throw more tantrums, a typical manifestation of mother-offspring conflict (Maestripieri, 2002). Maternal trade-offs over birth timing, which extend into mother-offspring conflict after birth, may commonly occur in long-lived species where development from birth to independence spans multiple seasons. Such trade-offs may substantially weaken the benefits of seasonal reproduction, and our findings therefore open new avenues to understanding the evolution of breeding phenology in long-lived animals, including humans.

**SIGNIFICANCE STATEMENT:** Why some species breed seasonally and others do not remain unclear. The fitness consequences of birth timing have traditionally been measured on offspring survival, ignoring other fitness components. We investigated the effects of birth timing on two fitness components in wild baboons, who breed year-round despite living in a seasonal savannah. Birth timing generates a trade-off between offspring survival and future maternal reproductive pace, meaning that mothers cannot maximize both. When birth timing favours maternal reproductive pace (instead of offspring survival), behavioural manifestations of mother-offspring conflict around weaning are intense. These results open new avenues to understand the evolution of reproductive timings in long-lived animals including humans, where such reproductive trade-offs may commonly weaken the intensity of reproductive seasonality.

## MAIN TEXT

## Introduction

Empirical studies investigating variation in reproductive timing have mostly focused on fast-lived seasonal breeders, whose development from birth to independence generally occurs within the most productive season (1). In long-lived mammals, the reproductive cycle from birth to weaning cannot similarly be squeezed into one annual food peak, and consequently, females must choose which stage(s) of the reproductive cycle to synchronize with one or more food peak(s). For example, female mammals could match the annual food peak to coincide with either late-weaning or mid-lactation, but usually not both. The reproductive timing strategy is likely to depend on how females trade-off the survival of their offspring (mortality risks tend to peak at the end of weaning) (2–4) with their own reproductive costs (energetic demands tend to peak around mid-lactation) (5, 6). Whether such reproductive timing strategies can vary within populations is largely unknown. In addition, while evolutionary trade-offs between offspring quality and quantity have been described both within and across species, through associations between birth spacing and infant growth and survival (7–11), the existence of maternal trade-offs over birth timing have only been suggested theoretically (3) and never tested empirically in mammals (but see for a bird species, *Fulica atra*: (12)).

Here, we investigate variation in maternal reproductive success and mother-offspring relationships associated with variable birth timings in the annual cycle of wild chacma baboons (*Papio ursinus*), living in a seasonal semi-arid savannah (Tsaobis, Namibia). Baboons are African primates distributed across a wide latitudinal range, and a classic model for understanding how early humans adapted to seasonal savannahs (13, 14). In particular, baboons typically breed year-round (15), and are therefore considered non-seasonal breeders, though the distribution of births shows moderate seasonality (i.e. varies along the annual cycle) in some species and populations (16–18). In addition, infant baboons, like many young primates including human toddlers, commonly perform tantrums, a manifestation of mother-offspring conflict (19–22). Using a combination of detailed long-term life-history and behavioural data collected over 15 years (2005-2019), we first characterize the reproductive and environmental seasonality of the Tsaobis baboons. Second, we quantify the consequences of birth timing on two components of female fitness: offspring survival and maternal inter-birth intervals (IBIs), and identify two distinct birth timing optima. We further test whether individual traits predict whether a female is more likely to give birth around one or the other optimum. Third, we investigate if compensatory maternal care mitigates the costs of suboptimal birth timing for offspring, and whether infants born, and subsequently weaned, in suboptimal timings increased their tantrum frequency.

## Results

### 1 Tsaobis baboons breed year-round despite living in a seasonal environment

Conceptions, births and cycle resumptions occurred throughout the year (Figure S1), indicating an absence of a strict breeding season. We used circular statistics to test whether moderate seasonality may still occur, computing respectively the mean annual angle (µ) and Rayleigh tests (R and p-values) for the annual distribution of 241 conceptions, 215 births and 171 cycle resumptions recorded between 2005-2019. The frequency of conceptions and births showed slight seasonal variations, which reached statistical significance for conceptions only (conceptions: µ = May 12, R=0.13, p=0.02; births: µ= November 18, R=0.09, p=0.17; cycle resumptions: µ= December 4, R=0.08, p=0.36, Figure S1).

Environmental seasonality was pronounced at Tsaobis (Figure 1A). Mean annual rainfall was low and variable (mean ± SD = 192 ± 143mm), falling mostly between January and April (Figure 1A). Following previous baboon studies (23, 24), we quantified food availability for baboons using the Normalized Difference Vegetation Index (NDVI), a satellite-based proxy of primary productivity with higher values corresponding to a higher degree of greenness (25). Seasonal variation in NDVI followed a similar, but slightly lagged pattern, to rainfall (Figure 1A). The highest birth frequency occurred in October-November, preceding the peak in rainfall (February) and NDVI (March-April, Figure 1A).

**Figure 1:**
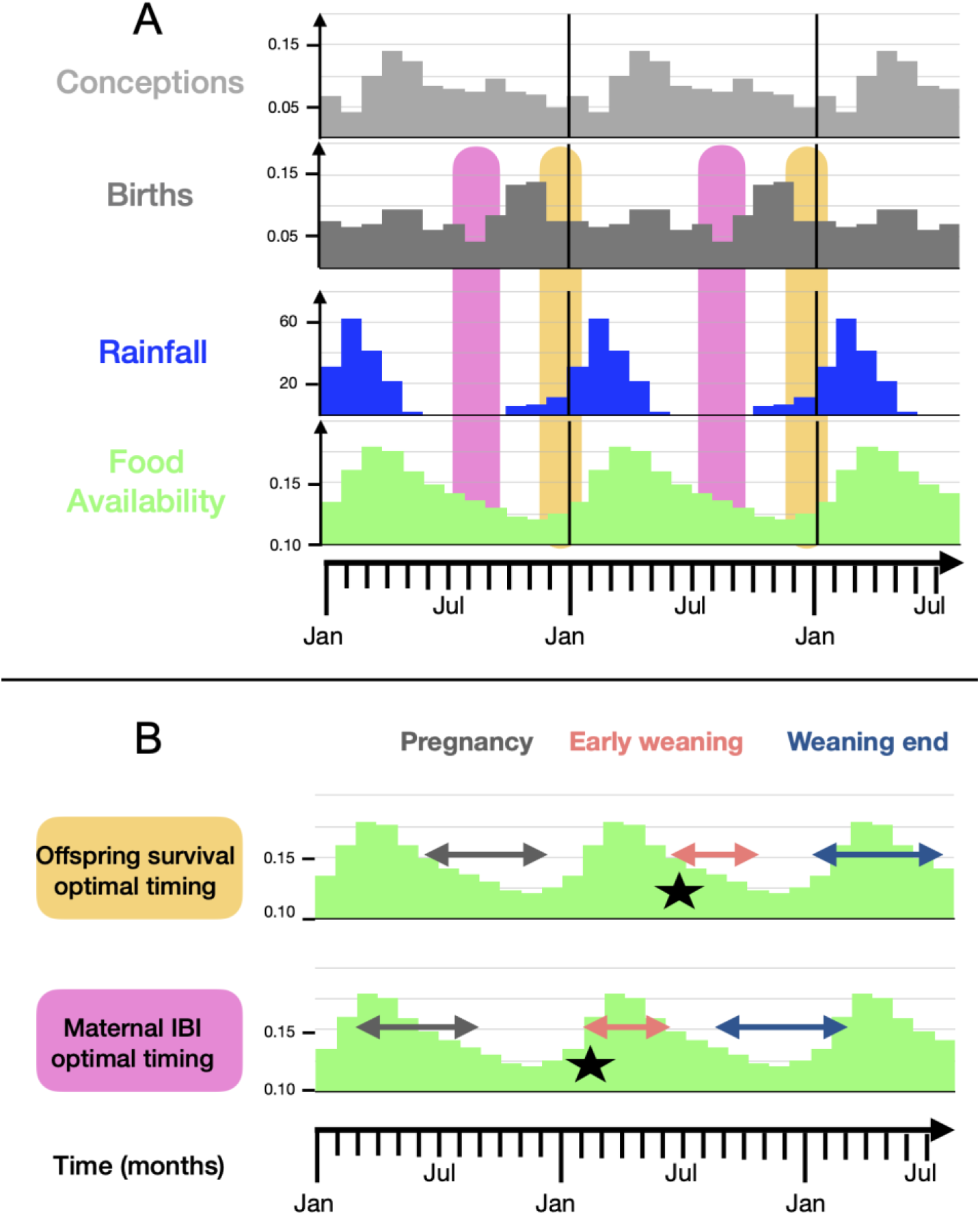
Tsaobis baboons’ reproductive timings in relation with environmental seasonality. In Panel A, we plotted the proportion of conceptions (N=241, in light grey) and births (N=215, in dark grey) recorded in 2005-2019 per month (Jan for January, Jul for July). We plotted the mean monthly cumulative rainfall (in mm) per month in blue and the mean NDVI value per month in green between 2005 and 2019. The orange and pink squares in the background represent resp. the offspring survival and the maternal IBI optimal birth timings. In Panel B, we aimed to represent the different phases of the female reproductive cycle, when the birth date occurs within the offspring survival (December 15^th^) or maternal IBI (September 1^st^) optimal timing, according to seasonal variation of NDVI. The green bar plot in the background indicates the mean NDVI per month (see y-axis). Pregnancy, indicated with grey arrows, occurs the 6 months prior a birth. Early-weaning, indicated with salmon-colour arrows, occurs from 6 to 9 months after a birth. Lactation peak, indicated with black stars, occur around 6 months after a birth. Weaning end, indicated with blue arrows, occurs from 12 to 18 months after a birth (see Appendix 4 for the characterization of these different reproductive stages).

### 2 Distinct birth timings optimize current versus future reproduction

There was an influence of birth timing on two indicators of maternal fitness. First, birth timing affects offspring survival (Table S1): infants born between November 15^th^ and January 1^st^ were the most likely to survive until weaning (Table S2), indicating an optimal birth timing for offspring survival in the annual cycle (Figure 2A). Second, the duration of maternal IBI is influenced by the timing of the birth opening the IBI (Table S1): females giving birth between August 1^st^ and September 15^th^ had the shortest IBIs (Table S2), indicating another different optimal birth timing for maternal reproductive pace in the annual cycle (Figure 2B). In the first case, the birth timing that maximises offspring survival synchronizes the seasonal food peak with the end of weaning, a highly vulnerable life stage for mammals (26–28), which occurs between 12 and 18 months after birth in this population (Figure 1B). In the second case, the birth timing that maximizes maternal reproductive pace synchronizes the food peak with the peak of lactation (occurring around 6 months after birth) (Figure 1B), which is the most energetically-costly reproductive stage for mothers (3, 26), and may therefore help to alleviate the costs of lactation and enhance maternal condition during the second half of lactation. The two months counting most births are October and November (i.e. 28.4% of annual births, Figure 1A) and the mean annual birth date is Nov 18^th^, indicating a reproductive trade-off over birth timing, that pushes mothers to target the very first days of the offspring survival optimal birth timing, in order to avoid compromising offspring mortality while minimizing the costs on their reproductive pace (Figure 2C).

**Figure 2:**
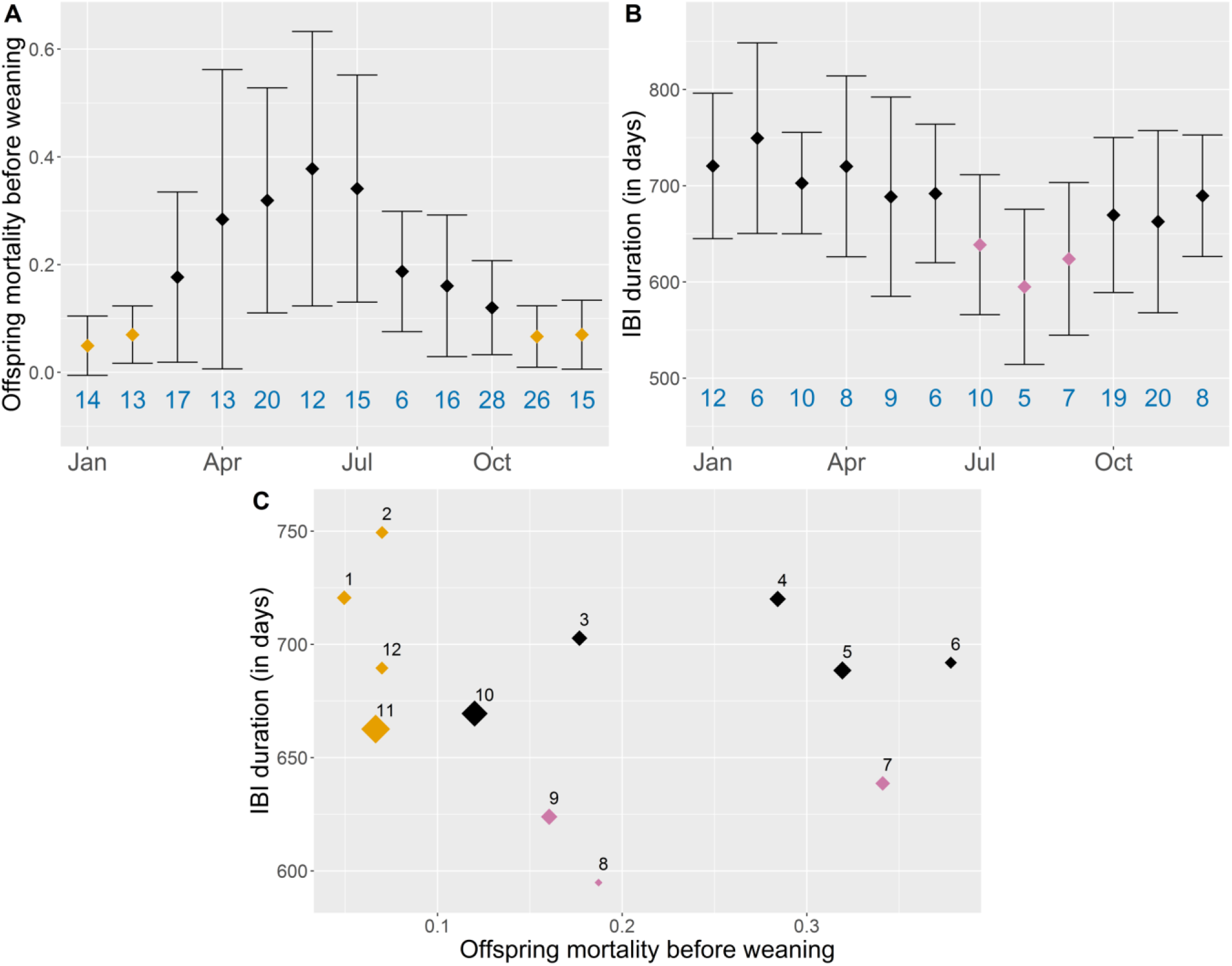
Distinct optimal birth timings for current and future reproduction. We plotted the predicted values of the full models (Model 1 looking at offspring mortality in panel A, and Model 2 looking at IBIs in panel B) according to the month of infant birth (Jan for January, Apr for April, etc.). The number of births observed for each month is indicated in blue below the bar. The dots represent the mean values, while the vertical black bars represent its standard deviations. The offspring survival optimal birth timing is identified as the period minimizing offspring mortality, i.e. from November to February, and indicated with orange dots (Panel A and C). The maternal IBI optimal birth timing is identified as the period minimizing maternal interbirth interval, i.e. from July to September, and indicated with pink dots (Panel B and C). In panel C, we represented the trade-off experienced by mothers over birth timing: each dot represents the predicted value of IBI according to the predicted value of offspring mortality for a given month birth (see label, 1 for January, 2 for February, etc.). The size of the dot is proportional to the number of births observed each month. We can notice an absence of points on the extreme bottom-left corner (i.e. with both low infant mortality - inferior to 0.10 - and short IBI - lower than 650 days), showing the existence of a reproductive trade-off. In addition, the highest number of births occur for the points closer to this bottom-left corner (months 10 & 11), indicating that mothers target a birth timing trading-off these two fitness components.

This result raises the possibility that some females might be more likely to time their births to maximise current over future reproduction, or vice versa. In particular, dominance rank and parity can affect various aspects of individual reproductive performance, including offspring survival and IBI (29–31), and may influence birth timing strategies accordingly. Similarly, mothers conceiving close to the optimal timing for maternal IBIs, which alleviates the energetic costs of lactation, may subsequently favour male over female embryos, which are more costly to produce in sexually dimorphic mammals (29, 32, 33). However, we failed to detect any significant variance associated with maternal identity on the deviation between observed birth and the optimal birth timing maximizing offspring survival (LRT=0.66, p=0.42) versus maternal IBI (LRT=0.00, p=0.98). This suggests that females did not consistently target one timing over the other across successive births. Moreover, female parity, rank and infant sex did not influence the proximity of birth timing in relation to each optimum (Table 1).

**Table 1:** Predictors of female reproductive timing. Estimates, confidence intervals, X^2^ statistics and P-values of the predictors of the two linear mixed models (Models 3 and 4). The response variables are respectively the deviation from the offspring survival optimal birth timing, i.e. from December 15^th^ (Model 3), and the deviation from the maternal IBI optimal birth timing, i.e. from September 1^st^ (Model 4), in days, based on 215 births from 62 females. Female identity and year of infant birth are included as random effects. For categorical predictors, the tested category is indicated between parentheses.

| Fixed effect |  | Estimate | IC |  | X<sup>2</sup> | P-value |
| --- | --- | --- | --- | --- | --- | --- |
|  |  |  | Lower | Upper |  |  |
| Model 3: Deviation from the offspring survival optimal birth timing |  |  |  |  |  |  |
| Infant sex | (Male) | 5.91 | -7.42 | 19.23 | 0.76 | 0.385 |
| Female parity | (Primiparous) | -12.77 | -30.02 | 4.47 | 2.11 | 0.147 |
| Female rank |  | 2.59 | -4.82 | 10.00 | 0.47 | 0.493 |
| Group | (L) | 5.16 | -10.37 | 20.69 | 1.33 | 0.515 |
|  | (M) | -12.66 | -44.70 | 19.39 |  |  |
| Model 4: Deviation from the maternal IBI optimal birth timing |  |  |  |  |  |  |
| Infant sex | (Male) | -3.19 | -16.46 | 10.08 | 0.22 | 0.7637 |
| Female parity | (Primiparous) | 9.67 | -7.49 | 26.82 | 1.22 | 0.269 |
| Female rank |  | -3.41 | -10.75 | 3.92 | 0.83 | 0.362 |
| Group | (L) | 10.67 | -4.70 | 26.04 | 1.92 | 0.382 |
|  | (M) | 0.93 | -30.91 | 32.78 |  |  |

### 3 Birth timings favouring future reproduction intensify mother-offspring conflict

In order to test whether maternal care may compensate for the costs of suboptimal birth timings in offspring, we investigated the effects of birth timing on the frequency of suckling and infant carrying. We did not find any effect of infant birth date on patterns of maternal care (Table S3). Further analyses revealed that mothers increase maternal care in the dryer winter months, but such compensation occurs regardless of an infant’s birth date (Appendix 1, Table S4).

We also investigated whether infants born in suboptimal timings may beg maternal care more frequently, looking at tantrum frequencies. We found that infants born near the maternal IBI optimal timing, i.e. between August 1^st^ and October 1^st^ (Table S2), were more likely to exhibit tantrums than other infants (Table S3, Figure 3). Observation date did not affect tantrum frequencies, meaning that such an effect was independent of the season of observation (Table S4).

**Figure 3:**
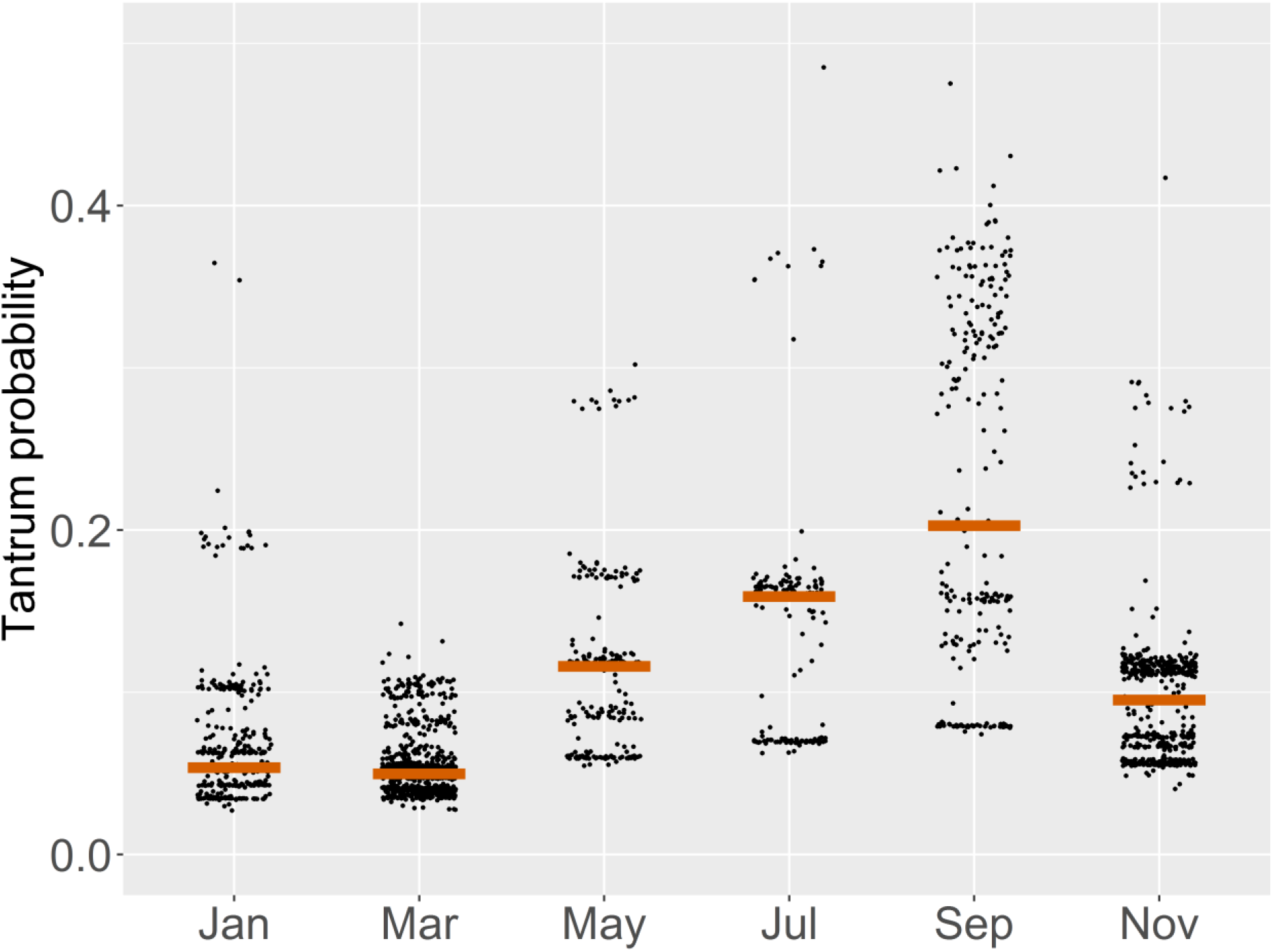
Influence of birth timing on tantrum probability. Predicted values of tantrum probability (Model 7) at weaning (age 12 months), according to infants’ birth month, based on 2221 focal observations from 55 infants. For graphical reasons, and given the low sample size of infants observed for some birth months, we pooled infants born in 2 consecutive months, so that Jan indicates infants born in both January and February, Mar in both March and April, etc. The brown horizontal bars indicate the median values of fitted values for each birth month category.

## Discussion

Our results further our understanding of the evolution of vertebrate reproductive timing in several ways. First, we identify two distinct optimal birth timings in the annual cycle, respectively favouring current reproduction (offspring survival) versus future reproduction (maternal reproductive pace). These are separated by four months, and the highest birth frequency occurs between these optima, indicating that mothers balance current and future reproduction, with a priority for offspring survival. Trade-offs over birth timing may be widespread in long-lived species with slow life histories, for which development from birth to independence spans several months, therefore exceeding the length of the most productive season. In such cases, different stage(s) of the reproductive cycle may be synchronized with one or more seasonal food peaks, with the specific pattern dependent on the trade-offs females make among different fitness components (34). Such variation could account for empirical cases where the observed birth peak fails to coincide with the birth timing expected on the basis of a single fitness measure. For example, in humans from pre-industrial Finland, births did not concentrate in the months with the highest infant survival expectations (35). More generally, such trade-offs may contribute to explain the partial or total lack of breeding seasonality observed in some large mammals (36), such as social primates including apes (18) and humans (37, 38).

Second, this study challenges the idea according to which non-seasonal breeding has evolved in response to an absence of optimal birth timing in the annual cycle, especially in species with ecological or physiological traits that buffer seasonal environmental variation (39). For example, chacma baboons and humans share a generalist diet (27), a capacity to extract fall-back foods at times of food scarcity (40), and an ability to store energy (2, 4), which have likely played a critical role in their adaptation to breed year-round in seasonal environments (41, 42). The few studies that have investigated the effects of birth timing on early survival of offspring in non-seasonal breeding primates such as geladas (*Theropithecus gelada*) (43) have indeed failed to detect any effect. In modern humans, fitness variation associated with seasonal birth timing is rare (38, 44), and where detectable, only has mild effects on adult longevity (45, 46). The fitness consequences of seasonal birth timing detected here were therefore unexpected, and surprisingly reveal that non-seasonal breeding can be favoured even where reproductive success depends on birth timing. Future work will usefully test the generality of these patterns in other species or populations that breed year-round to shed more light on the conditions favouring evolutionary transitions towards non-seasonal breeding.

Third, while different species synchronize different stages of their reproductive cycle with the seasonal food peak (1, 2, 47, 48), this study reveals variations in breeding timing within the same population. However, while mothers experience a trade-off between reproductive pace and offspring survival in their birth timing, it is not clear if particular individuals consistently favour certain strategies, as we did not detect any inter-individual effects of female identity, parity or rank on parturition timing. Instead, intra-individual factors, such as maternal reproductive history, may constrain the evolution of such individually-based specializations, if only because the duration of IBIs - 22 months on average but with extensive variation - prevents females from giving birth every two years at the same season. In addition, the costs of waiting for the next optimal timing may often outweigh the costs of giving birth at suboptimal timings. Overall, the fact that the highest frequency of births occur between the two optima or at the onset of the period maximizing offspring survival suggests that most females attempt to maximize offspring survival while optimizing their own reproductive pace.

Fourth, this study underlines the importance of weaning to understand the evolution of mammalian reproductive schedules. Late-weaning is most critical for infants who must learn to ensure their own provisioning. Matching that stage with the most productive season may substantially enhance infant survival (26, 27, 49, 50). Moreover, the peak of lactation typically coincides with the onset of weaning, and matching it with abundant resources can help to accelerate the transition to feeding independence by granting infants access to a wealth of weaning foods (Figure 1B) (26). Earlier weaning, in combination with better maternal nutritional condition, will likely promote the resumption of cycling (51–53), and may contribute to explain the shorter interbirth intervals associated with this birth timing. Such patterns may be very general. In the lemur radiation, for instance, despite a variety of life-histories, ecologies and societies, and the fact that different species mate and give birth at different times of year, all species synchronize weaning with the food peak (49). Our understanding of the ultimate causes of mammalian reproductive seasonality may gain from granting more consideration to the dynamics and consequences of weaning, which may have been underappreciated in comparison to the energetic costs of pregnancy and lactation (1, 2, 4).

Fifth, our results show that the trade-off over birth timing faced by mothers may subsequently translate into mother-offspring conflict after birth. Although mothers adjust maternal care seasonally, they do so regardless of the offspring’s age. Offspring born at suboptimal periods face the dry season in a critical developmental window (i.e., the end of weaning), and maternal care is insufficient to buffer them entirely from the adverse consequences that lead to higher mortality. Consequently, baboon infants respond by throwing more tantrums, which may be an honest signal of need (21, 54), just as children do in similar situations (22). More generally, these results shed light on the potential influence of environmental fluctuations, and specifically seasonality, on mother-offspring conflicts over maternal care. While the literature focusing on optimal birth spacing has mainly examined trade-offs between current and future reproduction for an implicitly stable level of resources (7, 55), such a stability may rarely be encountered by mothers in the wild (56, 57), who typically face extensive, but partly predictable, fluctuations in food availability. Taking into account the intensity and predictability of resource fluctuations may largely re-draw the landscape of strategic decisions available to mothers confronted with trade-offs between current and future reproduction in natural environments (57, 58).

Our findings open new perspectives to understand the evolutionary drivers of vertebrate reproductive seasonality, by revealing the occurrence of a maternal trade-off between current and future reproduction over birth timing, extended by mother-offspring conflict during weaning. Such a trade-off may commonly occur in organisms with a slow reproductive pace, and future studies on such taxa should investigate the consequences of reproductive timing on several fitness components. Indeed, multiple optimal birth timings in the annual cycle may generate a bimodal birth peak or an extended birth season. Evolutionary trade-offs over birth timing may therefore account for unexplained variation in the reproductive timing of long-lived vertebrates, including the evolution of non-seasonal breeding in humans and other species.

## Materials and methods

### Study site and population

Three habituated groups of wild chacma baboons were followed between 2005 and 2019: J and L since 2005, and M, a fission group from J, since 2016. They live in a desert-edge population at Tsaobis Nature Park (22°23S, 15°44’50E) in Namibia, in a strongly seasonal environment: the desert vegetation responds quickly to the austral summer rains, which usually fall between December and April, and then dies back during the dry winter months (59). Water is always available through the presence of both natural seeps and artificial water points for wildlife and livestock. A field team was present each year, mainly during winter (between May to October), for a variable number of months (mean = 4.5, range: 1.9-7.9), that collected daily demographic and behavioural data, as well as GPS locations, while following the groups on foot. All individuals, including infants, are individually recognizable thanks to small ear markings performed during capture and/or other distinctive features.

### Environmental data

In order to describe the relationship between reproductive and environmental seasonalities, we characterize two aspects of environmental seasonality at Tsaobis: rainfall and vegetation cover (an index of food availability).

Daily rainfall in a 0.25 × 0.25 degree grid cell resolution (corresponding to 28 × 28 km at this latitude) was extracted using satellite data sensors from the Giovanni NASA website (product TRMM 3B42) (60), from a rectangular geographic area encompassing the global ranging area of the Tsaobis baboons, computed using GPS locations collected by observers every 30 min when following the study groups. We used the minimal and maximal latitude and longitude recorded between 2005 and 2019. Monthly cumulative rainfall (summed across daily values) were computed between 2005 and 2019.

We used the Normalized Difference Vegetation Index (NDVI) as an index of food availability. NDVI is computed using the near-infrared and red light reflected by the surface of an area and measured with satellite sensors; it produces a quantitative index of primary productivity with higher values corresponding to a higher degree of vegetation cover (25). It has previously been used as an indicator of habitat quality for the Tsaobis baboons (24) and other baboon populations (23). We further confirmed that temporal variation in NDVI reflected temporal variation in rainfall: mean cumulative rainfall over the past three months explained between 60-72% of the NDVI variation (Appendix 2). To index food availability using NDVI for each troop, we first computed 100% isopleth home ranges for each group using kernel density estimates with the adehabitatHR package (‘kernelUD’ function) (61), based on the daily 30-min GPS locations from 2005-2019 (from 2016-2019 for M group). We obtained one home range per group for the entire study period. We then extracted the mean NDVI per 16 day-period on a 500 m × 500 m resolution across these periods using MODIS data (MODIS13A1 product) provided by NASA (25) within these home ranges for each group. Daily NDVI was computed by linear interpolation and then averaged to obtain a monthly value. In contrast to rainfall, NDVI was measured with greater resolution and for each group separately because baboons finely adjust their ranging behaviour in relation to food availability (62).

### Individual data

A female was considered adult when she reached menarche. The reproductive state of each adult female was monitored daily. A female could be: (i) pregnant (assessed by the paracallosal skin turning red and absence of cycles over the following months), with the exact start date of pregnancy being determined *post hoc* following infant birth, and encompassing 190 days (mean gestation length in this population, n = 13 pregnancies where both conception and birth were observed, range: 181-200 days, SD = 5) between conception and birth; (ii) lactating, as long as the female did not resume cycling after an infant birth; (iii) cycling, including both swollen females in oestrus (i.e., sexually receptive with a perineal swelling) and non-swollen females at other stages of their cycle. Conceptive cycles were established based on the beginning of a pregnancy, and were usually confirmed by a birth. The first post-partum cycle (i.e. cycle resumption) is the first cycle following an infant’s birth, when the female resumes cycling after lactation. The exact date of the cycle resumption corresponds to the first day of oestrus of the first post-partum cycle, i.e. the first day when a sexual swelling is recorded. The dates of these reproductive events (conceptions, births and cycling resumptions) were either known with accuracy when recorded by field observers, or estimated in the absence of observers using the methods detailed in Appendix 3 and Table S5.

Female parity was known from life history records and defined as primiparous (between the birth of her first and second infant) or multiparous (after the birth of her second infant).

Female social rank was established annually for each group using *ad libitum* and focal observations of agonistic interactions between adult females: supplants, displacements, attacks, chases and threats (Huchard and Cowlishaw 2011). We computed a linear hierarchy using Matman 1.1.4 (Noldus Information Technology, 2013), and then converted to a relative rank to control for group size (i.e. the number of adult females in the group). Each female was thus assigned one rank per year, ranging from 0 (lowest ranking) to 1 (highest ranking).

### Fitness data

We tested the influence of birth timing in the annual cycle on two fitness measures, namely offspring mortality before weaning and the duration of the maternal interbirth interval. For each infant born between 1^st^ January 2005 and 1^st^ August 2018, we investigated whether it died (yes/no) before weaning. The weaning age was identified as 550 days on the basis of the maximum length of post-partum anoestrus (n = 33 cases for which both birth and cycle resumption were known with accuracy, see also Appendix 4), and presumably reflected the upper threshold of weaning age in our population, assuming that females who resumed cycling had weaned their offspring, as lactation has suppressive effects on ovulation among primates (51, 52, 64). Death was recorded when a corpse was observed or when the infant had been missing in the group for five consecutive days. Infants born later than August 2018 were not considered as their survival outcome was unknown. Four infants that disappeared between consecutive field seasons were omitted because we could not establish whether the age of death was before or after 550 days. In our final dataset, a total of 39 infants out of 195 died before reaching 550 days of age, with mortality occurring at a median age of 74 days (range 1-284 days, n=17 known dates of death).

We defined interbirth intervals (IBI) as the number of days between two consecutive live births of the same female. We only considered IBIs for which the first infant reached weaning (29), i.e. survived until 550 days old. We discarded other IBIs as females resumed cycling rapidly after their infant’s death when unweaned (median=21 days, range=9-51, n=9 observed death), and their IBIs would have been shortened regardless of environmental seasonality. We computed a total of 120 interbirth intervals from 43 adult females, ranging from 397 to 1132 days with a mean of 678 days (SD=128).

### Behavioural observations

In order to characterize variation in maternal care and in mother-offspring conflict, we used three behavioural indicators: suckling, infant carrying and tantrum frequencies. We also used these behavioural data, along with life history data, to assign different developmental stages, including the different stages of weaning and the peak of lactation after an infant’s birth (see Appendix 4). In addition to life-history data, field observers collected behavioural data on infants aged between 2 and 24 months on a daily basis from dawn until dusk over four periods: from October to December 2006, from July to August 2017, from September to December 2018, and from April to July 2019. We collected a total of 1185 hours of focal observation on 69 infants across four field seasons (mean ± SD = 17.1 ± 7.8 hours of observations per infants, range = 6.3–34.6), with a mean of 40.7 focal observations per individual (SD=29.4). Focal observations were spread equally across the day (divided in four 3-h-long blocks) and focal individuals were chosen randomly, and never sampled more than once within a block. Focal observations durations were 1 h in 2006 and 20 min in 2017-2019, with a minimum of 10 min in all cases. We recorded the following activities on a continuous basis: suckling (when the focal individual had its mouth on its mother’s nipple; we could not distinguish comfort from nutritive suckling), travelling alone, infant carrying (carried by the mother, either ventrally or dorsally) and other activities. We also collected events related to mother-offspring conflicts (see below). In addition, we collected scan observations every 5 minutes (n=16702 scans across 3081 focal observations), including the activity of the focal individual.

#### Maternal care during weaning

Maternal care was quantified through two measures: suckling frequency and infant carrying frequency, which represent the two main energetic costs of maternal care before weaning (5, 65). First, for each scan observation (taken every 5 min), we considered whether the infant was suckling (1) or not (0) to investigate the effect of birth timing on variation in suckling frequency. In order to determine the best age window to consider, we explored age-related variation in suckling frequency, and found that suckling decreases gradually from 2 to 18 months old, before stabilizing to ca. 2% of the scans from 18 to 24 months old (Figure S2). In addition, the maximum length of post-partum amenorrhea, often used as a proxy for the end of weaning, lasted 550 days (i.e. 18.1 months) in this population (see above). Therefore, we considered only infants aged 2-to 18-months-old for this analysis, using 11687 scans from 55 infants. The birth date uncertainty for these 55 infants ranged from 0 to 130 days (with a median birth date uncertainty of 16 days) and was taken into account in subsequent models (see Appendix 5).

Second, for each scan observation during which an infant was travelling, we determined whether the infant was carried by its mother (1) or travelled on its own (0). This variable allowed us to monitor the gradual decrease from full maternal dependence to full independence during travelling. When looking for the best age window to consider, we observed that the proportion of infant carrying gradually decreases during the first year of life in our population (Figure S2), as in other baboon populations (65–67). Therefore, we considered infants aged from 2 to 12 months old for this analysis, using 924 scans from 35 infants.

#### Mother-infant conflicts during weaning

We considered infant tantrums as a behavioural measure of mother-offspring conflicts, reflecting when an infant’s request to access resources from its mother was not initially satisfied (19). Tantrum occurrence started in early-life, peaked when infants were aged around 6-9 months, and then gradually decreased during the end of their first and second year of life (Figure S2). Therefore, we considered only infants aged 2 to 18 months old for this analysis, using 2221 focal observations from 55 infants. During each focal observation, we determined if a tantrum occurred (1) or not (0), based on a range of distinctive offspring vocalizations (gecks, moans and loud screams) and behaviours (frenzied behaviour when infants hurl themselves to the ground, sometimes accompanied by rapidly rotating their tail) that were recorded on a continuous basis and are characteristic of baboon tantrums (21, 67). A tantrum was considered to occur when at least two of these behaviours or vocalizations were recorded, separated by at least 30s (isolated complaints, and complaints that lasted fewer than 30 seconds, were thus not considered as tantrums here). Tantrums were usually caused by maternal refusal of access to the nipple or to carrying, and more rarely by maternal absence.

### Statistical analysis

#### Characterization of the reproductive seasonality of the Tsaobis baboons

First, to assess the strength and direction of reproductive seasonality, we used a Rayleigh test, from circular statistics, which characterizes the deviation of circular data from a uniform distribution, via the mean direction µ and length R of the vector summing all observed events across the annual cycle (R=0 when the event is evenly distributed, and R=1 when all events are synchronized, i.e. occurs the same day) (68). Our sample comprised 241 conceptions, 215 births and 171 cycle resumptions which occurred between 2005 and 2019. Uncertainties in those dates were taken into account using 1000 randomized reproductive events for each variable (Appendix 5).

#### Birth timing effects on two fitness traits

To quantify the effect of birth timing on the probability of offspring mortality before weaning (Model 1), we ran a generalized linear mixed model (GLMM) with a binomial error structure. We then ran a linear mixed model (LMM, Model 2), testing the effect of birth timing on IBIs.

In both models, we used a sine term to describe the timing of an infant’s birth in the annual cycle. Sine waves allow the introduction of a circular variable into a multivariate model as a fixed effect: the possible effects of the date of birth are circular with a period of one year, as January 1^st^ is equally close to December 31^st^ than to January 2^nd^. This sinusoidal term was as follows:

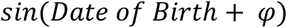

The date of birth in the formula above was converted in a radian measure, so that the period, i.e. one year, equalled to 2*π, ranging from 2*π/365 for January 1^st^ to 2*π for December 31^st^. We tested 12 different phase values *φ*(0, π/12, 2*π/12, 3*π/12, 4*π/12, 5*π/12, 6*π/12, 7*π/12, 8*π/12, 9*π/12, 10*π/12, 11*π/12), to account for different potential optimal periods for the event of interest across the year (24), as offspring mortality and IBI could be minimized for different birth dates (and so tested all potential dates as minimal). For example, a phase of 0 could maximize April 1^st^ or October 1^st^ depending on the sign of the estimate (see Table S2). We ran sequentially these 12 multivariate models, containing all other fixed and random effects (see below), and selected the best phase as the one minimizing the Akaike Information Criterion (AIC) in this full model set: the phase of 7*π/12 was retained for offspring mortality probability, and of 2*π/12 for IBI (Table S2). We controlled for birth date uncertainty in both models using a set of 1000 randomized birth dates within the interval of uncertainty (see Appendix 5 for more details).

In both models, we included as random effects year of infant birth and identity of the mother to control for repeated observations. In both models, we also included maternal parity, rank (in the birth year of the focal infant) and infant sex as fixed effects, because maternal parity and rank often affect reproductive traits in primates, including baboons (29, 69, 70), while infant sex can affect both the mother’s subsequent interbirth interval (71) and the probability of infant survival in sexually dimorphic primates (26, 32). We also control for group identity as a fixed effect in both models, as data were collected from only three groups in this study (72).

#### Characterization of optimal birth timings, and individual effects on birth timing

We investigated the individual determinants of female reproductive decisions over birth timing, based on 215 births from 62 females. We first used the results of Models 1 and 2 to characterize the optimal birth timings for offspring survival and maternal IBI respectively. Offspring mortality is minimised on December 15^th^ (Table S2), and we thus computed, for each birth date, the deviation in days from December 15^th^ (the maximum value of the deviation being 182 for June 15^th^). We used this deviation as a response variable of an LMM (Model 3) to investigate the individual determinants of giving birth close to, or away from, the timing that maximizes offspring survival. IBIs are minimised on September 1^st^ (Table S2), and we thus computed, for each birth date, the deviation in days from September 1^st^ (the maximum value of the deviation being 182 for March 1^st^). We used this deviation as a response variable of an LMM (Model 4), to investigate the individual determinants of giving birth close to, or away from, the timing that minimizes the maternal IBI.

For both Models 3 and 4, we tested the influence of infant sex, female parity and rank (as fixed effects) on the proximity of birth to the optimal timing for offspring survival (Model 3) or for maternal IBI (Model 4). We also controlled for the identity of the mother and birth year as random effects, and for group identity as fixed effects (as there was only three levels for this factor (72)). We tested the significance of maternal identity using a likelihood-ratio test (LRT), comparing the model with and without this random effect. We controlled for birth date uncertainty in these models using a randomization procedure described in Appendix 5.

#### Birth timing effects on maternal care and tantrum probability

We ran three GLMMs with a binomial error structure to test the effect of birth timing on the probability of suckling (Model 5), infant carrying (Model 6), and tantrums (Model 7). Models 5 and 6 are scan-based data: during a scan observation, the infant is suckling (yes/no, Model 5), and during a travelling scan observation, the infant is carried by its mother (yes/no, Model 6). Model 7 is based on the entire focal observation as tantrum events are relatively rare: during a focal observation, the infant throws a tantrum (yes/no).

In order to investigate the potential effect of birth timing on maternal care and tantrum probability, we used a sine wave term for infant birth date as a fixed effect, following the method explained for Models 1-2. We did so to examine natural minimums or maximums in the frequency of each of these traits along the annual cycle, without making any *a priori* hypothesis on which periods were minimized or maximized, in order to test whether the observed maximums or minimums would match the periods previously identified as the optimal timings favouring current versus future reproduction. We therefore tested 12 different phases in each full model and retained a phase of 9*π/12 for suckling, 0 for infant carrying, and 2*π/12 for tantrum probabilities. We controlled for birth dates uncertainty in all three models using the randomization procedure described in Appendix 5.

We included, as random effects, the identity of the infant (Models 5-7) to control for repeated observations. We also added the focal observation as a random effect for Models 5-6. We controlled for group identity and year of observation as fixed effects in all models, as there were less than five levels for both factors (72). In all models, we included maternal parity, rank (in the year of birth of the focal infant) and infant sex as fixed effects. Such parameters are likely to affect reproductive performances as well as the probabilities of maternal care and mother-offspring conflict (26, 67). For Model 7, we also controlled for the duration of focal observation as a fixed effect.

For Models 5-7, we further controlled for the effects of infant age, which modulates the amount of maternal care and probability of tantrums throughout early development (19, 26). We considered four different possibilities for the form of the relationship between infant age and the response variable, using a regression thin plate spline (general additive model), a simple linear effect, and a polynomial regression (of 2 or 3 degrees), respectively (73). To determine the best fit, we ran these different preliminary models with no other fixed effect but including all random effects (and the duration of focal observation for Model 7), and selected the model minimizing the AIC. The age effect was linear for suckling and infant carrying probabilities (Model 5 and 6), and a second-degree polynomial for tantrum probability (Model 7).

Lastly, mothers might be expected to invest more, and similarly infants might be expected to have more requests for maternal care, during the lean season, irrespective of the developmental trajectory of the infant, i.e. regardless of its age and birth timing (whether it was born in the optimal period or not). Therefore, we also investigated the potential effect of seasonality by assessing the influence of the observation date on suckling, infant carrying and tantrum probabilities (see Appendix 1 for more details). We did not include in the same model observation date and birth date, as they give redundant information (observation date is, by definition, the sum of birth date and infant’s age, and infant’s age is already included as a fixed effect). We present our models of birth date effects in the main text (Models 5-7, see also Table S3), and our models of observation date effects in the Supplementary Information (Models 5bis-7bis, Table S4).

The structure of each model, with the different fixed and random effects included, alongside sample size, is summarised in Table S6.

#### Statistical methods

All statistical analyses were conducted in R version 3.5.0 (74). For the Rayleigh test, we used the function ‘r.test’ from the R package ‘CircStats’ (75). To run mixed models, we used ‘lmer’ (for LMMs) or ‘glmer’ (for binomial GLMMs) function on the lme4 package (76). To run general additive mixed models (GAMMs) when investigating the best age effects on suckling, infant carrying and tantrum probabilities, we used the ‘gam’ function of the ‘mgcv’ package (73). All quantitative fixed effects were z-transformed to facilitate model convergence. When we obtained singular fits, we confirmed the results by running the same models with a Bayesian approach, using the ‘bglmer’ and ‘blmer’ functions of the ‘blme’ package (77). To diagnose the presence of multicollinearity, we calculated the variance inflation factor for each predictor in each full model using the ‘vif ‘function of the R ‘car’ package (78). These were lower than 2.5 in all cases. To assess the strength of the fixed effects in each model, we used the Wald chi-square tests with associated P-values computed with the ‘Anova’ function of the R package ‘car’ (78), and calculated the 95% Wald level confidence intervals. We further checked the distribution of residuals with ‘qqPlot’ function of the car package for LMMs (78), and with ‘simulateResiduals’ from DHARMa package for binomial GLMMs (79).

## DATA AND CODE AVAILABILITY

The datasets necessary to run analyses included in this paper and the associated legends have been deposited in the public depository: https://github.com/JulesDezeure/Maternal-trade-off-over-birth-timing-in-baboon

## ACKNOWLEDGMENTS

The authors are grateful to the Tsaobis Baboon Project volunteers from 2005 to 2019, and particularly to Harrison Anton, Charlotte Bright, Anna Cryer, Rémi Emeriau, Richard Gallagher, Chloe Hartland, Rachel Heaphy, Nick Matthews, Tess Nicholls, Vittoria Roatti and Ndapandula Shihepo for their dedicated effort at collecting focal observations on infant baboons. This research was carried out with the permission of the Ministry of Environment and Tourism, the Ministry of Land Reform, and the National Commission on Research, Science, and Technology. We further thank the Tsaobis beneficiaries for permission to work at Tsaobis, the Gobabeb Namib Research Institute and Training Centre for affiliation, and Johan Venter and the Snyman and Wittreich families for permission to work on their land. Data used in this study are part of long-term data collected within the framework of the Tsaobis Baboon Project, recently funded by a grant from the Agence Nationale de la Recherche (ANR ERS-17-CE02-0008, 2018-2021) awarded to EH. This paper is a publication of the ZSL Institute of Zoology’s Tsaobis Baboon Project. Contribution ISEM n°XX.

