## Supplementary Material for "Breeding seasonality generates reproductive trade-offs in a long-lived mammal"

### SUPPLEMENTARY TEXT

**Appendix 1.** Seasonal effects on maternal care and tantrum probability

When modelling suckling, infant carrying and tantrum probabilities (Models 5-7), we further tested for seasonal effects, i.e. effects of the date of observation, on the response variable. To do so, we applied the approach used to describe the effects of birth timings, i.e. a sine term of the date of observation (in radians) was entered as a fixed effect in the multivariate model. This sinusoidal term was as follows:

$$sin(Date\ of\ observation + \varphi)$$

The date of observation in the formula above was converted to a radian measure, so that the period, i.e. one year, equalled 2*π, ranging from 2*π/365 for the 1st of January to 2*π for the 31st of December. We similarly tested 12 different phase values $\varphi$ and selected the best phase as the one minimizing the AIC of the full models 5-7 (with all random and fixed effects, except the sine term of the date of birth). We found that 7*π/12 was the best phase for suckling probability, 3*π/12 for infant carrying probability, and 10*π/11 for tantrum probability. The results of the models with observation date are presented in Table S4.

**Appendix 2.** Correlations between rainfall and NDVI at Tsaobis

In order to estimate the correlation between monthly rainfall and NDVI at Tsaobis, we used a moving window approach. We expected that the cumulative rainfall over the preceding months, rather than the rainfall during the current month, would be the best predictor of monthly NDVI. First, we identified the time window maximizing the correlation between rainfall and NDVI variation, testing periods covering 0 to 6 month(s) prior to the current month using an AIC-based selection procedure, and a univariate linear model containing only the fixed effect of interest (cumulative rainfall over variable periods) and three response variables, namely the monthly NDVI values associated with the home ranges of all three study groups. For these three groups, the time window minimizing the model AIC was cumulative rainfall over the preceding three months. The adjusted value of the model $R^2$ measures the proportion of NDVI variance explained by variation in cumulative rainfall over the past three months.

**Appendix 3.** Estimations of the dates of conceptions, births and cycle resumptions

We characterized the reproductive seasonality in our population considering three main reproductive events: conceptions, births, and cycle resumptions (i.e. the end of post-partum amenorrhea).

1. ***Births.*** The dates of births, conceptions, and cycle resumptions were directly observed where possible, but otherwise estimated for those periods when no observers were present. We observed a total of 84 births. Of those, 62 were seen by observers on the exact day, and 22 were witnessed after a short absence (leading to a small uncertainty in the actual date: median=17 days, range=1-30). When the conception only was observed (n=52 births), we estimated birth dates by adding the mean gestation length (n=190 days, range: 181-200 days, SD=5, n=13 pregnancies where both conception and birth were observed) to the conception date. Conception was considered to occur on the day of deturgescence (D-day) of the swelling during a conceptive cycle. This generated a total of 136 birth dates known with high accuracy. When neither conception nor birth were observed (n=56 births), the birth date was estimated using infant coloration (based on the progressive loss of natal coat and skin coloration) following a method recently described and validated in our population (1), with further refinement provided by the reproductive history of the mother (e.g., if the mother was pregnant during the last three months of a field season, then the infant was necessarily born in the three months following the end of this season given that a pregnancy lasts 190 days). Finally, when neither birth nor conception was observed, and infant colour when first seen was unknown or uninformative (i.e.,

the transition from natal to adult coat had already occurred) (n=23 births), we used the reproductive state of females in the preceding field season to minimize uncertainty over birth timing. For example, if a female was cycling the last day of the preceding season, the infant was necessarily born at least 190 days after this day. In total, our sample comprised 215 births between 2005 and 2019, with a median uncertainty of 10 days (range: 0-153 days) (Table S5).

2. ***Conceptions.*** We observed 81 conceptions: 68 conceptions were witnessed (observers were present during the conceptive cycle), and 13 occurred during a short absence of observers (leading to a small uncertainty in the actual date: median=10 days, range=3-30). The exact date of conceptions was the day of swelling detumescence of the conceptive cycle (when witnessed) (2). When the birth was observed but not the conception, the latter was estimated to occur 190 days prior to birth (n=65 conceptions). When neither conception nor birth were observed but birth occurred (i.e. no miscarriage), we estimated birth date as explained above, and inferred conception from the birth date using the mean gestation period (n=79). Finally, when pregnancy signs were seen (i.e. red paracallosal skin and an absence of cycles) but conception was not observed and there was no birth due to a miscarriage or early death (occurring before an infant was recorded by observers), we estimated the date of conception using female reproductive states (n=16 conceptions). For example, if a female was seen pregnant on the first day of a field season, we knew that the conception occurred from 0-190 days prior to this date (as gestation lasts 190 days in this population). All in all, we generated a sample of 241 conceptions, with a median uncertainty of 10 days (range: 0-164 days).

3. ***Cycle resumptions.*** Only cycle resumptions following a period of lactation were included in our analyses. Cycle resumptions were observed in 64 cases. In 107 other cases, a female was lactating at the end of a field season and then cycling at the beginning of the next (median days between consecutive field seasons=225, range=83-584). To reduce the

uncertainty of the resumption date estimations in these cases, we calculated the minimum post-partum amenorrhea length (the time between birth and cycle resumption) (mean ± SD = 353 ± 89 days, range=223-550) based on the 33 cases for which both events were known, and used this value as a minimum threshold in our estimations. We also excluded all cycle resumptions for which the uncertainty exceeded one year. Our sample thus comprised a total of 171 cycle resumptions, with a median uncertainty of 61 days (range= 0-272 days).

**Appendix 4.** Characterization of developmental stages: weaning and lactation peak

In order to understand which stage of the reproductive cycle might be timed with the seasonal food peak (Figure 1B), we needed to define the sequential behavioural stages of weaning, which is the infant's gradual transition to nutritional independence (3), along with the peak of lactation in our population. First, the onset of weaning can be defined as the period when solid foods start to account for an important part of an infant's energy intake, and is characterized by an increase in maternal refusals to accede to her offspring's suckling demands. The onset of behavioural conflict between a mother and her offspring has therefore often been used as a proximate measure of the early-weaning period (3). In our population, tantrum probabilities peak between 6 and 9 months of age (Figure S2), and we therefore used this age window to characterize 'early-weaning'. The peak of lactation typically occurs just before the beginning of weaning (3, 4), when offspring have grown larger but are still fully dependent. So we can consider that lactation peak occurs around 6 months after birth in our population. Similar ages for early-weaning and lactation peak have been found in other baboon populations (5–7), albeit weaning age and lactation durations can vary substantially between populations (8).

The end of weaning can be defined as the complete cessation of nursing, i.e. when offspring feed exclusively on solid foods. Looking at behavioural data, suckling frequencies decrease gradually from 2 to 18 months old, before stabilizing to ca. 2% of time from 18 to 24 months old (Figure S2). In addition, the maximum length of post-partum amenorrhea (PPA), often used as a proxy for the end of weaning (3), was 550 days (i.e. 18.1 months) in this population (based on n = 33 cases for which both birth and cycle resumption were known with accuracy). We therefore considered the age of 18.1 months as an upper threshold marking the end of weaning for all juveniles in our models on infant mortality and IBI (see main text). However, age at the end of weaning is highly variable between individuals, as indicated by the wide range of variation for PPA (8-18 months, mean = 12 months). To take this variation into account, we defined the 'end of weaning' as the age window of 12-18 months after birth for Figure 1B.

All in all, for Figure 1B, in order to better understand the relationship between reproductive phenology and environmental seasonality, we considered the lactation peak to occur around 6 months after birth, early-weaning between 6 and 9 months of age, and the end of weaning between 12 and 18 months of age. As a note of caution, these windows are strictly based on behavioural and life-history data, which show some limitations to evaluate the dynamics of lactation (9, 10). More objective measures, such as isotopic comparisons of mother-offspring hair or faecal samples (11–14), may help to refine these estimates.

**Appendix 5.** Controlling for uncertainties in the dates of conceptions, births and cycle resumptions in statistical analyses

Dates of conceptions, births and cycle resumptions were estimated in many cases because the Tsaobis baboons are not followed year round (see Appendix 1). In addition, uncertainty in these estimations varied with the time of year, as we generally follow baboons during the cooler, dryer months. In order to account for these uncertainties in our analyses, we ran a set of randomizations to evaluate the robustness of the fixed effects found to be statistically significant. For each reproductive event (conceptions, births, and cycle resumptions) for which the date was associated to some uncertainty (i.e. exact date unknown), we created an extended dataset including all possible dates of the full range of uncertainty (from the minimum to the maximum date). For example if a baboon birth date was estimated to occur between October 2$^{nd}$ and December 23$^{rd}$, we included all possible dates between October 2$^{nd}$ and December 23$^{rd}$ in this extended dataset. Using this extended dataset, we generated 1000 simulations; in each iteration, a date was randomly drawn between the minimal and maximal estimate for each reproductive event that was not known with certainty. Events known with certainty did not vary throughout such simulations.

These simulations were integrated in different statistical analyses slightly differently. In our characterization of reproductive seasonality, we extracted the mean R, μ and p-value of the Rayleigh test for the 1000 simulated datasets of cycle resumptions, conceptions and births. We indicate these mean values in the main text. We also computed the 95% level confidence intervals of these p-values: for conceptions, p=0.019 – 0.021; for births, p=0.166 – 0.174; for cycle resumptions, p=0.328 – 0.358.

In our multivariate mixed models investigating the effect of seasonal birth timing on offspring mortality before weaning (Model 1) and maternal interbirth intervals (Model 2), we controlled for the uncertainties in dates of birth which could affect both our response variables and our main fixed effect of interest (seasonal birth timing). For Model 1, we generated 1000 simulations with random birth dates drawn, for each birth, between minimal and maximal birth date estimations for this particular birth, and subsequently ran 1000 mixed models, one for each simulated value of the offspring's birth date and for each survival outcome (as birth date affects an offspring's age, and therefore its estimated age at death). For Model 2, we similarly generated 1000 simulations with random birth dates drawn between minimum and maximum birth date estimations for the two births defining the IBI. We subsequently ran 1000 models with randomized values for both IBI (the response variable, i.e. number of days between the first and second birth) and the birth date fixed effect. For both models, we then extracted the 1000 p-values of our fixed effect ‘seasonal birth timing’ and computed the confidence intervals of these p-values (see the footnote of Table S1).

In our analysis of the individual determinants of birth timing (Models 3 and 4), we similarly generated 1000 simulations of birth dates drawn between minimum and maximum birth date estimations, and assessed for each of these randomly drawn births the deviation, in days, from December 15th for Model 3 and September 1st for Model 4 respectively. For both models, we then ran 1000 LMMs using these deviations as the response variable (Models 3 and 4). We extracted 1000 p-values of our various fixed effects, and computed their 95% level confidence intervals. No fixed effect was close to significance, and this information was thus not added to the footnote of Table 1.

Finally, in our analysis investigating the effects of seasonal birth timing on maternal care and mother-offspring conflict at the behavioural level, we similarly generated 1000 simulations of birth dates drawn between minimum and maximum birth date estimations. We then ran 1000 GLMMs looking at the effect of seasonal birth timing, along with other covariates, on the probabilities of suckling (Model 5), infant carrying (Model 6) and tantrums (Model 7). We extracted the 1000 resulting p-values for our fixed effect ‘seasonal birth timing’, computed their median and 95% level confidence interval, and added this information to the footnote of Table S3.

### TABLES

**Table S1**: Predictors of offspring mortality before weaning and maternal interbirth interval (IBI) duration.

Estimates, confidence intervals, $X^2$ statistics and P-values of the predictors of a binomial generalized mixed model of the probability of offspring mortality before weaning (0/1: survived/died, Model 1) and a linear mixed model of the duration of the maternal birth interval (IBI) (in days, Model 2), based on 195 observations from 57 females for Model 1 and 120 observations from 43 females for Model 2. Female identity and year of infant's birth are included as random effects in both models. Significant effects are indicated in bold. For the fixed effect 'birth date', we also indicate in the footnote the 95% confidence interval of the average p-value of the simulated models taking into account birth date uncertainty. Infant birth date is fitted using a sine term with a phase of 7*π/12 for infant mortality and of 2*π/12 for IBI, and. For categorical predictors, the tested category is indicated between parentheses.

| Fixed Effect | | Estimate | IC | | $X^2$ | P-value |
|---|---|---|---|---|---|---|
| | | | Lower | Upper | | |
| **Model 1: Offspring mortality** | | | | | | |
| **Infant birth date** | | **-1.12** | **-1.84** | **-0.40** | **9.38** | **0.002*** |
| Infant sex | (Male) | 0.20 | -0.76 | 1.15 | 0.16 | 0.685 |
| Female parity | (Primiparous) | -0.83 | -2.32 | 0.67 | 1.17 | 0.279 |
| Female rank | | -0.44 | -0.95 | 0.07 | 2.87 | 0.090 |
| Group | (L) | -1.29 | -2.39 | -0.18 | 5.25 | 0.072 |
| | (M) | -0.10 | -4.08 | 3.88 | | |
| **Model 2: Maternal IBI** | | | | | | |
| **Infant birth date** | | **36.84** | **7.59** | **66.09** | **6.10** | **0.014 †** |
| Infant sex | (Male) | 37.04 | -0.36 | 74.45 | 3.77 | 0.052 |
| Female parity | (Primiparous) | 44.53 | -3.24 | 92.29 | 3.34 | 0.068 |
| **Female rank** | | **-25.73** | **-50.68** | **-0.77** | **4.08** | **0.043** |
| Group | (L) | **-50.41** | -105.19 | 4.38 | 3.31 | 0.191 |
| | (M) | -31.09 | -150.57 | 88.38 | | |

* 95% CI: [0.00967 – 0.01087]

† 95% CI: [0.02533 – 0.02821]

**Table S2:** Identification of the best birth timing effect for Models 1-2 & 5-7: ΔAIC (Akaike Information Criterion) according to the phase of the sine wave term

In order to identify the best birth timing effect on our various indicators of fitness and maternal care, we ran 12 different models, with 12 different phases $\varphi$ for the sine wave term of the birth date (as a fixed effect), for each full model (Models 1-2 & 5-7). If the estimate of the sine term fixed effect is positive, then the birth date maximised is indicated in the 'Date maximised' column and the one minimised is indicated in the 'Date minimised' column. On the contrary, if the estimate of the sine term fixed effect is negative, then the birth date maximised is indicated in the 'Date minimised' column. ΔAIC of each model equals the AIC value of the considered model minus the AIC value of the best model (ΔAIC=0 for the best model, indicated in bold writing). We selected the best phase as the one minimizing the AIC, i.e. for which ΔAIC=0. For example, for Model 2, the best phase is $\varphi = 2 * \pi/12$, and the estimate of the sine term fixed effect is positive (Table S1), indicating that IBIs are maximised in March 1st, and minimized in September 1st. Wherever the fixed effect 'birth date' was significant (Model 1, 2 and 7), we considered all phases $\varphi$ for which ΔAIC<2 to define the optimal time window presented in the main text (see greyer background), for instance between August 1st and September 15th for Model 2.

| Phase $\varphi$ | Date maximised | Date minimised | Δ AIC | | | | |
|---|---|---|---|---|---|---|---|
| | | | Mortality (Model 1) | IBI (Model 2) | Suckling (Model 5) | Infant carrying (Model 6) | Tantrum (Model 7) |
| 0 | 1st April | 1st October | 8.26 | 2.16 | 0.67 | **0** | 1.15 |
| π/12 | 15th March | 15th September | 9.42 | 0.72 | 1.13 | 0.17 | 0.34 |
| 2*π/12 | 1st March | 1st September | 9.45 | **0** | 1.62 | 0.65 | **0** |
| 3*π/12 | 14th February | 15th August | 8.12 | 0.33 | 1.96 | 1.29 | 0.39 |
| 4*π/12 | 1st February | 1st August | 5.66 | 1.61 | 1.92 | 1.85 | 1.49 |
| 5*π/12 | 15th January | 15th July | 2.91 | 3.36 | 1.46 | 2.16 | 2.94 |
| 6*π/12 | 1st January | 1st July | 0.84 | 5.02 | 0.86 | 2.18 | 4.20 |
| 7*π/12 | 15th December | 15th June | **0** | 6.16 | 0.37 | 1.93 | 4.84 |
| 8*π/12 | 1st December | 1th June | 0.47 | 6.59 | 0.09 | 1.53 | 4.80 |
| 9*π/12 | 15th November | 15th May | 1.99 | 6.29 | **0** | 1.05 | 4.20 |

| | | | | | | | |
|---|---|---|---|---|---|---|---|
| 10*π/12 | 1st November | 1st May | 4.10 | 5.31 | 0.08 | 0.56 | 3.27 |
| 11*π/12 | 15th October | 15th April | 6.34 | 3.84 | 0.32 | 0.17 | 2.19 |

**Table S3:** Birth timing and other predictors of the probability of suckling, infant carrying and tantrums

Estimates, confidence intervals, $X^2$ statistics and P-values of the predictors of binomial generalized linear mixed models of the probability of suckling (Model 5), infant carrying (Model 6) and tantrums (Model 7), including infant's identity (and focal number for Models 5 and 6) as random effect, and focal observation time as an 'offset' fixed effect for Model 7. These GLMMs are based on 11687 scan observations from 55 infants for Model 5, 924 scan observations from 35 infants for Model 6 and 2211 focal observations from 55 infants for Model 7. Significant effects are indicated in bold. For relevant significant effect, we also indicated in the footnote the 95% level confidence interval of the 1000 p-values taking into account birth date uncertainty. Infant birth date is fitted as a sine term with a phase of 9*π/12 for suckling, 0 for infant carrying, and 2*π/12 for tantrum probabilities. For categorical predictors, the tested category is indicated between parentheses.

| Fixed Effect | | Estimate | IC Lower | IC Upper | $X^2$ | P-value |
|---|---|---|---|---|---|---|
| **Model 5: Suckling** | | | | | | |
| Infant birth date | | -0.32 | -0.76 | 0.13 | 1.97 | 0.16 |
| Infant sex | (Male) | -0.01 | -0.73 | 0.73 | 0.00 | 0.99 |
| Female parity | (Primiparous) | -0.89 | -2.20 | 0.42 | 1.76 | 0.18 |
| Female rank | | -0.09 | -0.42 | 0.24 | 0.30 | 0.59 |
| **Infant age** | | **-1.66** | **-1.97** | **-1.35** | **110** | **<$10^{-4}$** |
| **Group** | **(L)** | **0.18** | **-0.53** | **0.90** | **7.79** | **0.02** |
| | **(M)** | **1.51** | **0.36** | **2.66** | | |
| **Observation year** | **(2017)** | **0.21** | **-1.22** | **1.63** | **40.45** | **<$10^{-4}$** |
| | **(2018)** | **1.73** | **0.84** | **2.62** | | |
| | **(2019)** | **0.03** | **-0.82** | **0.88** | | |
| **Model 6: Infant carrying** | | | | | | |
| Infant birth date | | 0.53 | -0.18 | 1.24 | 2.13 | 0.14 |
| **Infant sex** | **(Male)** | **-0.94** | **-1.60** | **-0.28** | **7.78** | **0.005** |
| **Female parity** | **(Primiparous)** | **-1.08** | **-2.11** | **-0.05** | **4.21** | **0.040** |
| **Female rank** | | **-0.51** | **-0.82** | **-0.19** | **10.0** | **0.002** |
| **Infant age** | | **-1.94** | **-2.69** | **-1.20** | **26.2** | <$10^{-4}$ |

| | | | | | | |
|---|---|---|---|---|---|---|
| Group | (L) | 0.01 | -0.53 | 0.55 | 0.67 | 0.71 |
| | (M) | -0.35 | -1.34 | 0.64 | | |
| Observation year | (2017) | -11.5 | -262 | 239 | 3.69 | 0.30 |
| | (2018) | 1.36 | -0.36 | 3.09 | | |
| | (2019) | 0.86 | -1.19 | 2.91 | | |
| **Model 7: Tantrum** | | | | | | |
| **Infant birth date** | | **-0.32** | **-0.62** | **-0.03** | **4.53** | **0.033*** |
| Infant sex | (Male) | -0.10 | -0.45 | 0.25 | 0.33 | 0.57 |
| Female parity | (Primiparous) | 0.35 | -0.32 | 1.02 | 1.02 | 0.31 |
| Female rank | | 0.03 | -0.14 | 0.19 | 0.11 | 0.74 |
| **Infant age** | **Age** | **-41.13** | **-54.06** | **-28.21** | **53.28** | **$<10^{-4}$** |
| | **$Age^2$** | **-19.07** | **-28.21** | **-9.92** | | |
| Group | (L) | -0.35 | -0.68 | -0.01 | 4.23 | 0.12 |
| | (M) | -0.21 | -0.81 | 0.40 | | |
| **Observation year** | **(2017)** | **0.12** | **-1.39** | **1.63** | **12.61** | **0.006** |
| | **(2018)** | **0.30** | **-0.81** | **1.41** | | |
| | **(2019)** | **-0.44** | **-1.57** | **0.68** | | |
| **Offset** | | **0.40** | **0.10** | **0.71** | **6.81** | **0.009** |

* 95% CI: [0.04684 – 0.05234]

**Table S4**: Seasonality and other predictors of the probability of suckling, infant carrying and tantrums

Estimates, confidence intervals, $X^2$ statistics and P-values of the predictors of the binomial GLMMs of the probability of suckling (Model 5bis), infant carrying (Model 6bis), and tantrums (Model 7bis). Each model includes infant's identity and year of infant's birth as random effects. Models 5bis and 6bis also included focal observation as random effects, whereas Model 7bis included focal observation time as an offset fixed effect. Observation date is fitted as a sine term with a phase of $7*\pi/12$ for suckling probability, $3*\pi/12$ for infant carrying probability, and $10*\pi/12$ for tantrum probability. Significant effects are indicated in bold. For categorical predictors, the tested category is indicated between parentheses.

| Fixed Effect | | Estimate | IC | | $X^2$ | P-value |
|---|---|---|---|---|---|---|
| | | | Lower | Upper | | |
| **Model 5bis: Suckling** | | | | | | |
| **Observation date** | | **-1.66** | **-2.91** | **-0.40** | **6.70** | **0.0096** |
| Infant sex | (Male) | -0.03 | -0.79 | 0.74 | 0.00 | 0.95 |
| Female parity | (Primiparous) | -0.89 | -2.25 | 0.47 | 1.63 | 0.20 |
| Female rank | | -0.11 | -0.46 | 0.23 | 0.40 | 0.53 |
| **Infant age** | | **-1.62** | **-1.94** | **-1.30** | **97.47** | **<10-4** |
| **Group** | **(L)** | **0.30** | **-0.46** | **1.06** | **6.71** | **0.035** |
| | **(M)** | **1.52** | **0.33** | **2.71** | | |
| **Observation year** | **(2017)** | **-2.34** | **-4.88** | **0.20** | **29.99** | **<10-4** |
| | **(2018)** | **1.74** | **0.82** | **2.66** | | |
| | **(2019)** | **-2.64** | **-4.95** | **-0.33** | | |
| **Model 6bis: Infant carrying** | | | | | | |
| **Observation date** | | **-1.14** | **-1.98** | **-0.30** | **7.12** | **0.0076** |
| **Infant sex** | **(Male)** | **-0.90** | **-1.56** | **-0.24** | **7.20** | **0.0073** |
| Female parity | (Primiparous) | -0.78 | -1.77 | 0.22 | 2.33 | 0.13 |
| **Female rank** | | **-0.45** | **-0.76** | **-0.15** | **8.31** | **0.0039** |
| **Infant age** | | **-2.40** | **-2.83** | **-1.97** | **120** | **<10-4** |
| Group | (L) | -0.003 | -0.53 | 0.54 | 2.04 | 0.36 |
| | (M) | -0.63 | -1.62 | 0.37 | | |
| **Observation year** | **(2017)** | **-13.10** | **-2410** | **2384** | **12.2** | **0.007** |
| | **(2018)** | **0.66** | **-1.19** | **2.52** | | |
| | **(2019)** | **-0.73** | **-2.77** | **1.31** | | |
| **Model 7bis: Tantrum** | | | | | | |
| Observation date | | 0.63 | -0.16 | 1.42 | 2.44 | 0.12 |
| Infant sex | (Male) | -0.09 | -0.45 | 0.26 | 0.27 | 0.60 |
| Female parity | (Primiparous) | 0.07 | -0.57 | 0.72 | 0.05 | 0.83 |
| Female rank | | 0.04 | -0.13 | 0.21 | 0.23 | 0.63 |
| **Infant age** | **Age** | **-33.10** | **-43.45** | **-22.75** | **51.09** | **<10-4** |
| | **$Age^2$** | **-20.84** | **-29.87** | **-11.81** | | |

| | | | | | | |
|---|---|---|---|---|---|---|
| Group | (L) | -0.32 | -0.66 | 0.02 | 3.86 | 0.15 |
| | (M) | -0.05 | -0.64 | 0.53 | | |
| Observation year | (2017) | 0.57 | -1.12 | 2.27 | 0.45 | 0.93 |
| | (2018) | 0.20 | -0.91 | 1.30 | | |
| | (2019) | 0.45 | -1.18 | 2.09 | | |
| **Offset** | | **0.40** | **0.10** | **0.71** | **6.73** | **0.0095** |

**Table S5**: Different methods used to estimate the dates of births of the 215 baboon infants born at Tsaobis between 2005 and 2019.

| Criteria used for estimation | Infant colour when first seen | N births estimated | Median uncertainty (days) | Range of uncertainty (days) |
|---|---|---|---|---|
| Birth observed in the field | Pink | 62 | 0 | 0 |
| Birth occurred during a short field break | Pink | 22 | 17 | 1-30 |
| Conception date known | / | 52 | 10 | 10- 37 |
| Infant coloration & mother's reproductive state (1) | Pink or transitional | 56 | 61 | 6-151 |
| Mother's reproductive state only | Grey or unknown | 23 | 67 | 21-153 |
| Total | / | 215 | 30 | 0-153 |

**Table S6**: Summary of the structure of all models included in the study.

| Indicators | Fitness traits | | Birth timing | | Maternal care | | |
|---|---|---|---|---|---|---|---|
| Model number | 1 | 2 | 3 | 4 | 5 | 6 | 7 |
| Response variable | Offspring survival before weaning | Interbirth intervals (days) | Deviation from the offspring survival optimal birth timing | Deviation from the maternal IBI optimal birth timing | Suckling | Infant carrying | Tantrum |
| Model type | Binomial GLMM | LMM | LMM | LMM | Binomial GLMM | Binomial GLMM | Binomial GLMM |
| Number of observations | 195 | 120 | 215 | 215 | 5089 | 924 | 2221 |
| Number of individuals (juveniles / mothers) | 57 | 43 | 62 | 62 | 55 | 35 | 55 |
| Fixed effects | Infant birth date, infant sex, female parity, female rank, group | Infant birth date, infant sex, female parity, female rank, group | Infant sex, female parity, female rank, group | Infant sex, female parity, female rank, group | Infant birth date (or observation date, see Table S4), infant sex, female parity, female rank, infant age, group, observation year | Infant birth date (or observation date, see Table S4), infant sex, female parity, female rank, infant age, group, observation year | Infant birth date, (or observation date, see Table S4), infant sex, female parity, female rank, Infant age$^2$, group, observation year, focal duration |
| Random effects | Birth year, female identity | Birth year, female identity | Birth year, female identity | Birth year, female identity | Infant identity, focal number | Infant identity, focal number | Infant identity |

FIGURES

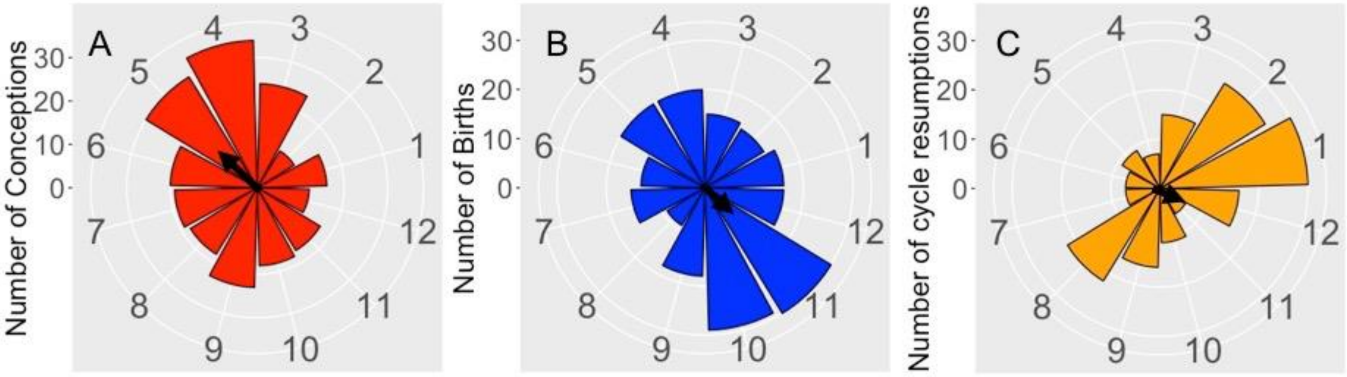

**Figure S1**: Chacma baboons breed all year round.

Number of conceptions (Panel A, N=241), births (Panel B, N=215) and cycle resumptions (Panel C, N=171) per month (from 1=January to 12=December) between 2005 and 2019. Births and cycle resumptions do not show significant seasonality, while conceptions significantly deviate from non-seasonality, with an average conception date in May. The black arrow length is the value of the Rayleigh statistic R, and its direction is μ. The numbers on the y-axis of each plot indicate the scale for the number of events on that plot.

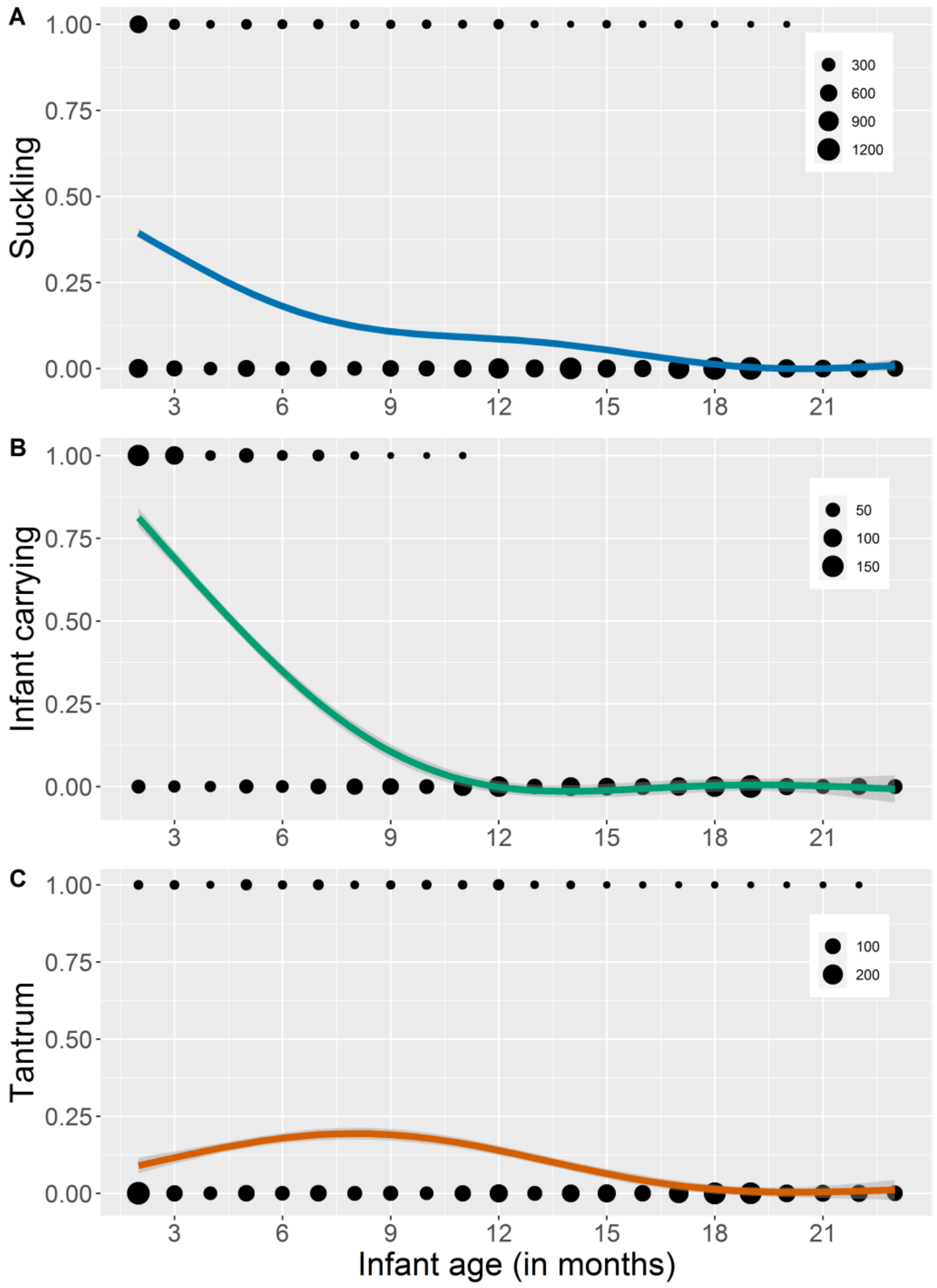

**Figure S2:** Variation in the probabilities of suckling, infant carrying, and tantrums, according to infant age.

We plotted (A) the probability of suckling during a scan, (B) the probability of infant carrying during a travelling scan, and (C) the probability of tantrum during a focal observation according to infant age (in months). For all panels, the size of black dots is proportional to the number of observations (see plot legends). The coloured curves show the predicted fit using a general additive function (method 'gam' of geom_smooth function in 'ggplot2' R package). The darker area around each curve represents the confidence interval of the fitted curve.
